# Neutrophils Suppress Mucosal-Associated Invariant T Cells

**DOI:** 10.1101/745414

**Authors:** M Schneider, RF Hannaway, R Lamichhane, SM de la Harpe, JDA Tyndall, AJ Vernall, A Kettle, JE Ussher

## Abstract

Mucosal-associated invariant T (MAIT) cells are innate-like T lymphocytes that are abundant in mucosal tissues and the liver where they can respond rapidly to a broad range of riboflavin producing bacterial and fungal pathogens. Neutrophils, which are recruited early to sites of infection, play a non-redundant role in pathogen clearance and are crucial for controlling infection. The interaction of these two cell types is poorly studied. Here, we investigated both the effect of neutrophils on MAIT cell activation and the effect of activated MAIT cells on neutrophils. We show that neutrophils suppress the activation of MAIT cells by a cell-contact and H_2_O_2_ dependent mechanism. Moreover, highly activated MAIT cells were able to produce high levels of TNFα that induced neutrophil death. We therefore provide evidence for a negative regulatory feedback mechanism in which neutrophils prevent over-activation of MAIT cells and, in turn, MAIT cells limit neutrophil survival.

## Introduction

Mucosal-associated invariant T (MAIT) cells are innate-like T cells that are rapidly activated by a broad range of bacterial and fungal pathogens.^1^ MAIT cells are characterised by a semi-invariant T cell receptor (TCR), Vα7.2-Jα12/20/33, and are restricted by the MHC class I-related protein 1, MR1.^2^ Their activating ligand is derived from 5-amino-6-D-ribitylaminouracil (5-A-RU), an intermediate in the riboflavin synthesis pathway, which is common in bacteria and fungi, but absent in humans.^3, 4^

MAIT cells represent the most abundant innate-like T cell subset in human blood, comprising approximately 5% of T cells, and are enriched at mucosal sites, including the lungs and liver.^1^ MAIT cells are rapidly activated in response to riboflavin-producing bacteria via a TCR-dependent mechanism, producing effector cytokines (interferon-γ (IFNγ), tumour necrosis factor α (TNFα), and interleukin-17 (IL-17)), showing cytotoxic potential (through release of perforin and granzymes), and undergoing proliferation.^5, 6, 78^ TCR-independent activation by cytokine stimulation, including by IL-12 and IL-18, results in a comparatively delayed response and altered effector responses.^6, 8^

MAIT cells have been shown to influence various immune cell subsets to promote activation of the adaptive immune response. In a mouse model of *Francisella tularensis* pulmonary infection, monocyte differentiation to monocyte-derived dendritic cells was driven by the MR1-independent production of GM-CSF by MAIT cells, affecting recruitment and activation of CD4^+^ T cells.^9, 10^ MAIT cells can furthermore instruct dendritic cell (DC) maturation *in vitro* through a CD40L and MR1-dependent mechanism, promoting IL-12 production.^11^ Additionally, in a TCR-dependent manner, MAIT cells can provide help to B cells, inducing plasmablast differentiation and antibody production *ex vivo*.^*12*^ Overall, alongside their cytotoxic effector functions, MAIT cells appear to have important immune modulatory functions, particularly for regulation of myeloid cells.

Neutrophils are the most numerous myeloid cell type in the blood and play a crucial role in pathogen clearance.^13, 14, 15^ They are rapidly recruited to sites of inflammation.^15^ MAIT cells are able to produce IL-17A, which can promote the production of the neutrophil chemokine CXCL8.^7, 16, 17^ In addition, activated MAIT cells can directly induce neutrophil chemotaxis.^8^ Therefore, MAIT cells may play a role in the recruitment of neutrophils to the site of infection. However, little is known about the interaction between neutrophils and MAIT cells at the site of infection. Neutrophils have been shown to suppress activation of conventional T cells and innate-like T cells, including γδ T cells and invariant natural killer T (iNKT) cells,^18, 19, 20, 21^ however, their effect on MAIT cell activation is yet to be explored. Similarly, the effect of activated MAIT cells on neutrophils has been sparsely studied. Davey *et al*. showed that both Vδ2^+^ γδ T cells and MAIT cells had a protective effect on neutrophil survival and induced an antigen presenting phenotype in neutrophils, dependent on GM-CSF, IFNγ, and TNFα.^22^

Here, we set out to investigate the relationship between neutrophils and MAIT cells in the presence of a bacterial stimulus. We aimed to understand how neutrophils influence MAIT cell activation and, in turn, how ligand activated MAIT cells affect neutrophils to better understand their roles in response to bacterial infection.

## Results

### Neutrophils are poor activators of MAIT cells and suppress MAIT cell activation

To assess if neutrophils are able to activate MAIT cells in response to bacteria, neutrophils and Vα7.2^+^ cells (referred to as MAIT cells) were incubated with fixed *E*. *coli* for 6 or 22 hours. MAIT cell activation was assessed by intracellular cytokine staining for IFNγ and TNFα (Figure 1A). Neutrophils were unable to stimulate significant IFNγ production by MAIT cells at either 6 or 22 hours, or TNFα production at 6 hours (Supplementary figure 1A-C). However, a small population of MAIT cells were stimulated to produce TNFα at 22 hours, which was dependent on MR1 but not IL-12 (Figure 1B). To investigate whether the presence of another antigen presenting cell may enhance activation, CD14^+^ monocytes were added to the co-culture. MAIT cells were activated well in the presence of CD14^+^ monocytes alone, however this activation was significantly reduced with addition of neutrophils and did not differ significantly from MAIT cell activation in the presence of neutrophils alone (Figure 1C+D). Consistent with the effect observed with *E*. *coli* treatment, neutrophils alone were poor activators of MAIT cells with 5-A-RU treatment and suppressed MAIT cell activation by monocytes (Figure 1E+F). Little IFNγ production was seen with 5-A-RU treatment, but a similar proportion of MAIT cells produced TNFα in response to 5-A-RU as the co-cultures treated with *E*. *coli* (Figure 1E+F). Neutrophil mediated suppression of MAIT cell TNFα production was significant, but not IFNγ production.

**Figure 1:**
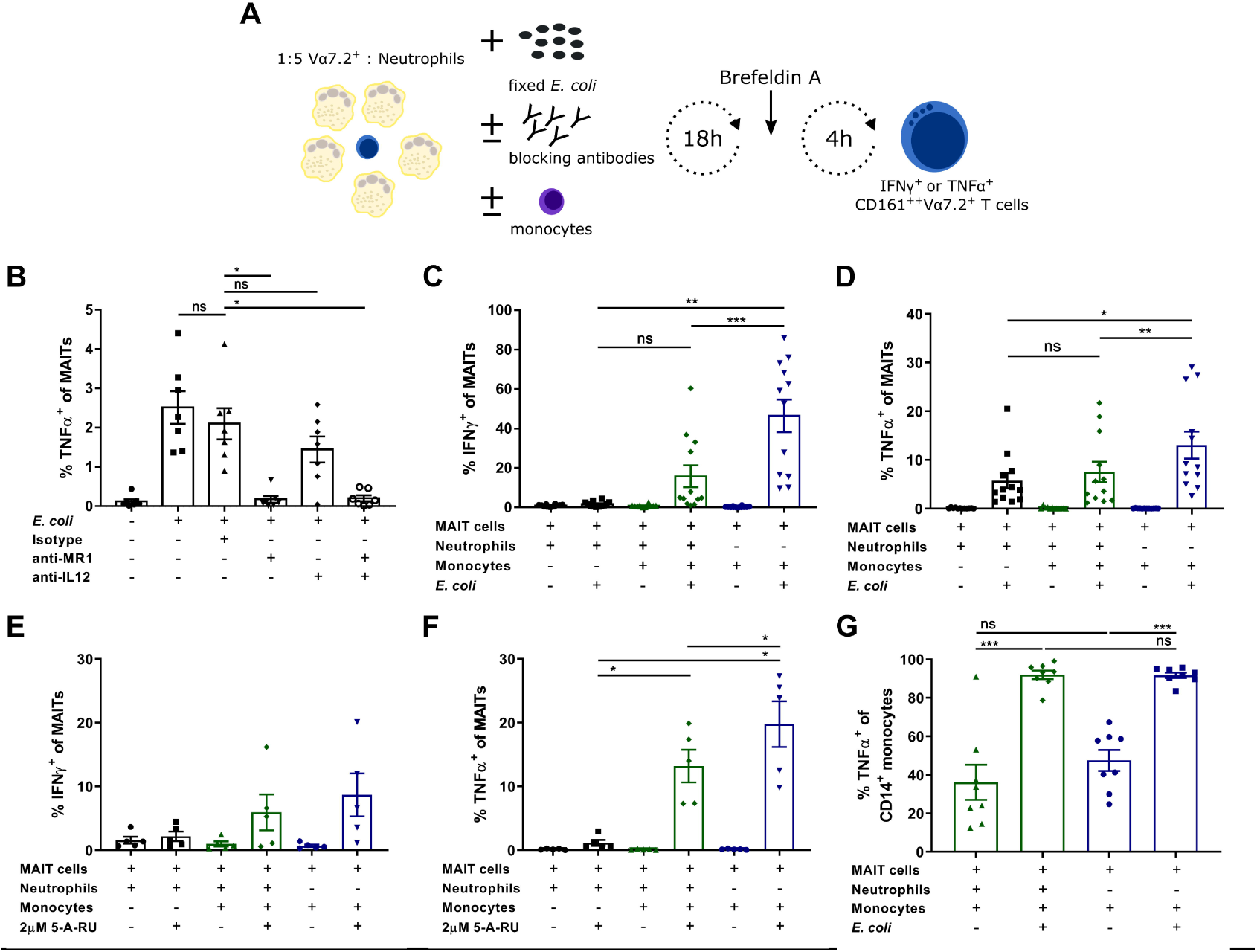
Neutrophils are poor activators of MAIT cells and inhibit MAIT cell activation by monocytes. **A** Vα7.2^+^ cells and neutrophils (1:5 ratio) were activated with 10bpc fixed *E*. *coli* (**B-D**, and **G**) or 2 µM 5-A-RU (*E-F*) and incubated *B* for 22 hours ± blocking antibodies against MR1 and/or IL-12, or *C-D* for 22 hours or *E-G* 6 hours ± CD14^+^ monocytes (1:1 ratio to Vα7.2^+^ cells). MAIT cell activation was assessed by intracellular staining for TNFα and IFNγ by flow cytometry. TNFα and/or IFNγ production in MAIT cells (CD161^++^Vα7.2^+^CD3^+^) is shown for **B** (n=7), **C-D** (n=12), **E-F** (n=5), and **G** (n=8). Graphs show mean with SEM. Each data point represents an individual blood donor. **B** Repeated measures one-way ANOVA with Tukey multiple comparisons test, or **C-G** Repeated measures one-way ANOVA with Bonferroni’s multiple comparison test of preselected pairs. * p<0.05, ** p<0.01, *** p<0.001, ns = not significant.

Analysis of monocyte activation, as assessed by TNFα production in CD14^+^ cells, showed no significant effect of neutrophils on monocyte activation. Treatment with fixed *E*. *coli* resulted in TNFα production in over 90% of monocytes after 6 hours of incubation, regardless of neutrophil presence or absence (Figure 1G). A trend towards suppression of activation was seen with neutrophil presence when mean fluorescence intensity (MFI) was assessed, however, this did not reach significance (Supplementary Figure 1D). Therefore, neutrophils suppress the activation of MAIT cells, but have no significant effect on CD14^+^ monocyte activation.

Next, we investigated whether this suppression of MAIT cells by neutrophils is similarly evident in whole PBMCs. As neutrophils have previously been shown to suppress γδ T cell activation, comparison was made to the effect of neutrophils on Vδ2^+^ γδ T cell (Vδ2^+^ T cell) activation.^19^ To ensure the effect seen was not as a result of bacterial phagocytosis by neutrophils, PBMCs were treated in one of two ways: cells were mixed with equal numbers of neutrophils and co-cultured (c) directly with fixed *E*. *coli* for 6 hours, or PBMCs were pre-incubated (p) with fixed *E*. *coli* for 20 min and then washed before neutrophil addition. MAIT cell and Vδ2^+^ T cell activation were analysed by flow cytometry for activation marker expression (Figure 2A). The presence of neutrophils significantly inhibited the activation of both MAIT cells and Vδ2^+^ T cells in PBMCs independently of neutrophil exposure to bacteria. MAIT cells and Vδ2^+^ T cells showed a marked decrease in TNFα and IFNγ production alongside reduced CD69 upregulation and, for MAIT cells but not Vδ2^+^ T cells, 4-1BB upregulation (Figure 2B+C). While degranulation, as measured by CD107a surface expression, was significantly suppressed in MAIT cells in the presence of neutrophils, no significant change was seen in the expression of granzyme B and perforin in either cell subset (Figure 2C). As previously reported, Vδ2^+^ T cells generally expressed higher levels of both granzyme B and perforin than MAIT cells.^23, 24, 25^ IL-17A production was insubstantial both in the absence and presence of neutrophils in both cell types (Figure 2C).

**Figure 2:**
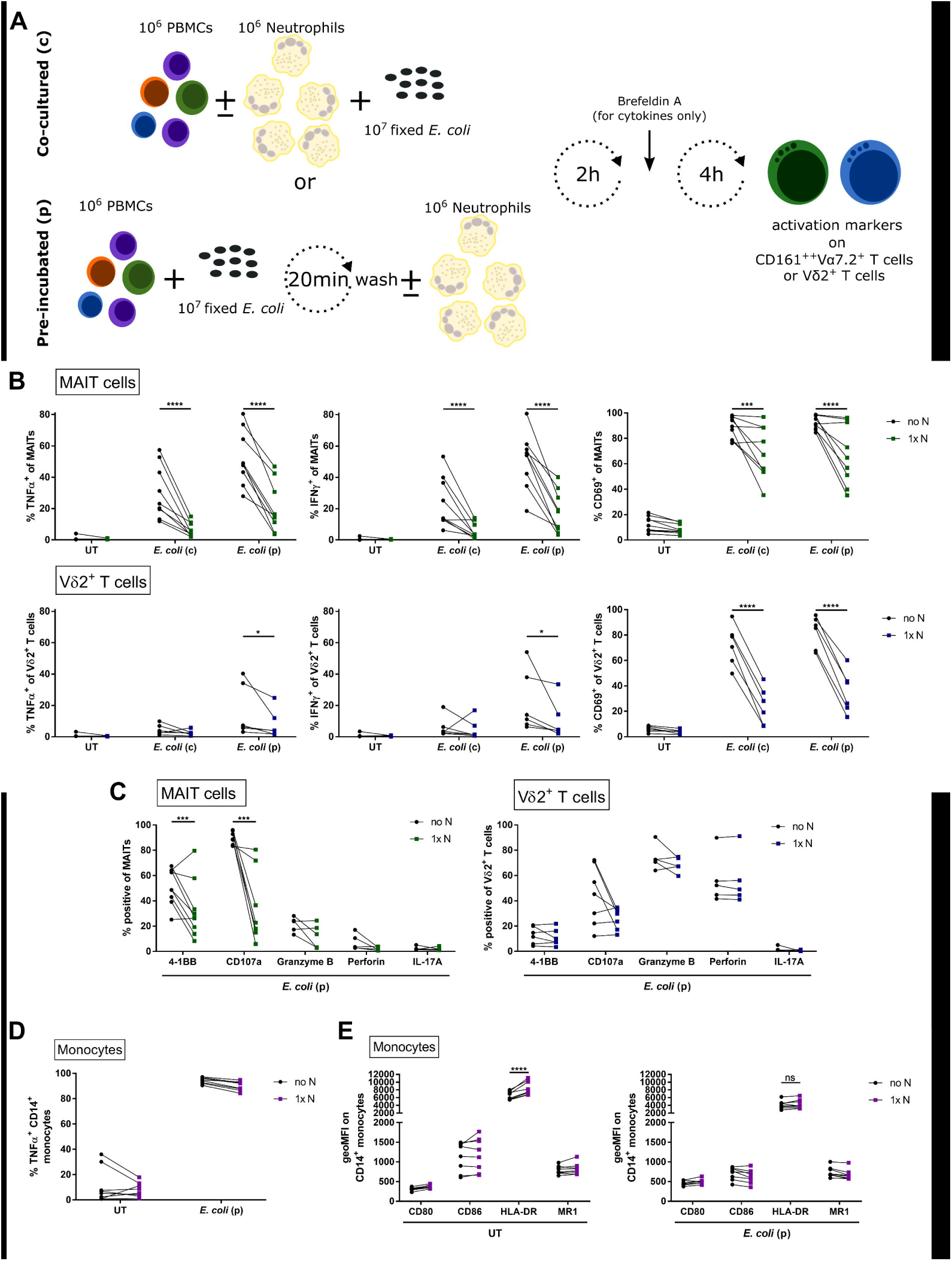
Neutrophils suppress MAIT and Vδ2^+^ T cell activation in PBMCs. **A** Equal numbers of freshly isolated PBMCs and neutrophils (1x N) were mixed and activated with 10bpc fixed *E*. *coli* either by adding bacteria to the co-culture (c) (**B**) or by pre-incubating (p) PBMCs with bacteria for 20 min and washing before addition of neutrophils (**B**-**E**). Control samples without addition of neutrophils (no N) were run in parallel. Samples were incubated for 6 hours, with brefeldin A added for the last 4 hours if cytokine production was assessed. Activation markers were analysed in MAIT cells (CD161^++^Vα7.2^+^CD3^+^) and Vδ2^+^ CD3^+^ T cells. **B** TNFα, IFNγ, and CD69 expression in MAIT cells (top) and Vδ2^+^ T cells (bottom) is shown (n=9 and n=6, respectively). **C** Expression of activation markers 4-1BB (n=9 for MAIT cells; n=6 for Vδ2^+^ T cells), CD107a (n=7), granzyme B, perforin, and IL-17A (all n=5) in MAIT cells and Vδ2^+^ T cells is presented. **D** TNFα expression (n=8) and **E** geometric mean of fluorescence intensity of CD80, CD86, HLA-DR, and MR1 on CD14^+^ monocytes is shown (n=6). **B**-**E** RM two-way ANOVA with Bonferroni’s multiple comparisons test was performed. Each data point represents an individual blood donor. * p<0.05, *** p<0.001, **** p<0.0001, ns = not significant.

The effect of neutrophils on monocyte activation in PBMCs was analysed by TNFα expression in CD14^+^ monocytes. Over 90% of monocytes expressed TNFα in response to *E*. *coli* treatment; this was not significantly affected by the presence of neutrophils (Figure 2D). However, when TNFα expression was assessed by MFI it was significantly reduced with neutrophil presence (Supplementary Figure 1E). Assessment of the effect of neutrophils on markers of activation and antigen presentation, including CD80, CD86, and MR1, on CD14^+^ monocytes in response to *E*. *coli* treatment showed no significant effect. However, HLA-DR expression on CD14^+^ monocytes was significantly increased in the presence of neutrophils in untreated, but not *E*. *coli* treated, samples (Figure 2E).

Overall, this data shows that neutrophils are poor activators of MAIT cells and have an inhibitory effect on the activation of MAIT cells and Vδ2^+^ T cells in the presence of other antigen presenting cells. This observed suppression appears to be a direct effect of neutrophils on MAIT cells and Vδ2^+^ T cells, as neutrophils had little effect on the activation or phenotype of CD14^+^ monocytes.

### Mechanism of suppression

Following the above findings, we went on to investigate the mechanism through which neutrophils suppress MAIT cells. Firstly, to determine if the inhibition of MAIT cell activation by neutrophils was dependent on neutrophil viability, we analysed the ability of fixed neutrophils to suppress the activation of MAIT cells. Fixed neutrophils were less able to inhibit MAIT cell activation, showing only a mild suppression of IFNγ production (Figure 3A). In contrast, Vδ2^+^ T cell activation was suppressed by fixed neutrophils equivalently to fresh neutrophils (Figure 3A). Next, to analyse the importance of cell contact on neutrophil inhibition of MAIT cell and Vδ2^+^ T cell activation, neutrophils and PBMCs were separated using transwell plates. Control plates lacking transwells were analysed in parallel. The inhibitory effect of neutrophils on MAIT cell and Vδ2^+^ T cell activation was abolished with neutrophil and PBMC separation; no reduction of TNFα and IFNγ production was seen in samples separated by transwells. The loss of this suppressive effect of neutrophils was not overcome by using three times the amount of neutrophils (Figure 3B).

**Figure 3:**
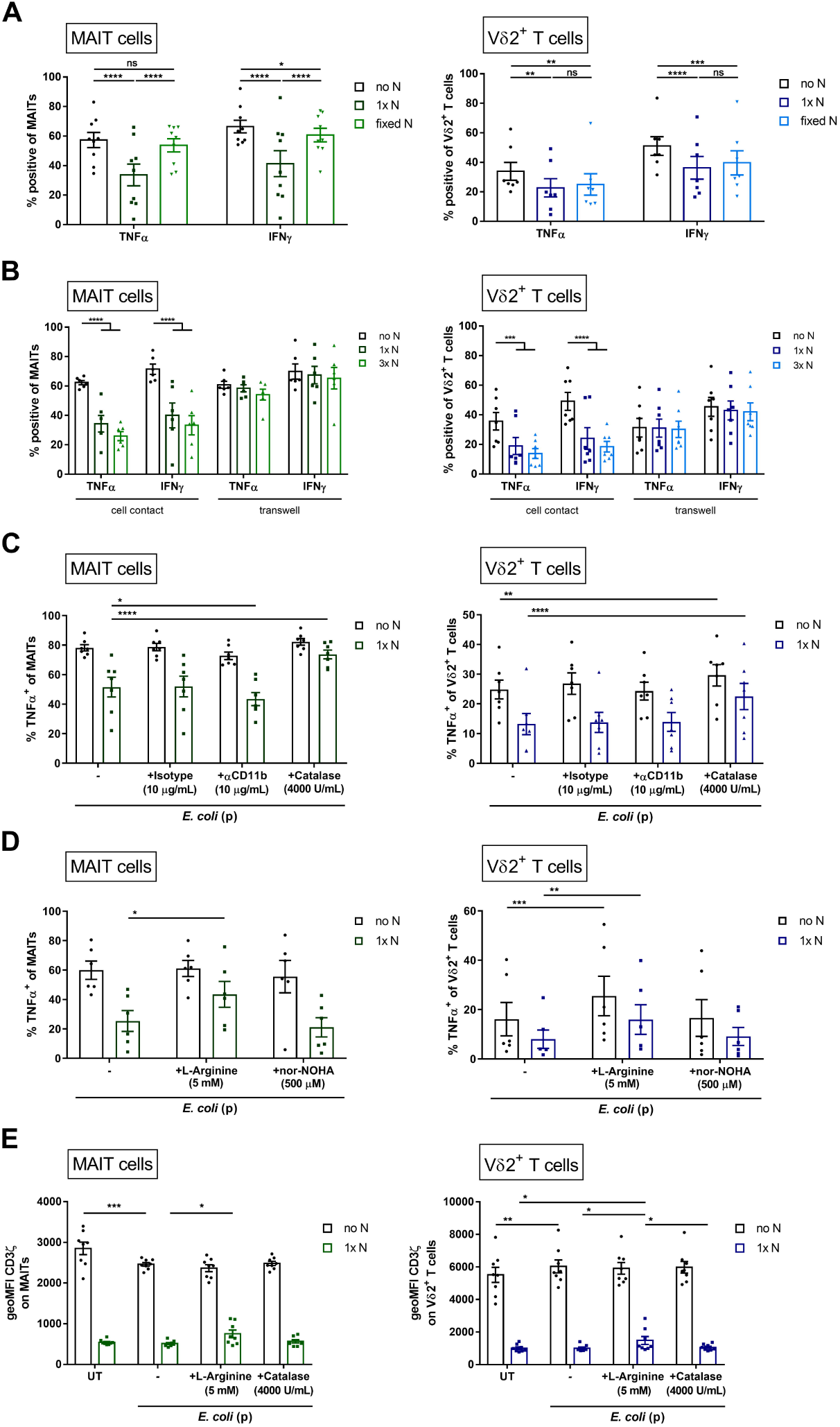
Neutrophils suppress MAIT cells and Vδ2^+^ T cells by a cell-contact and H_2_O_2_ dependent mechanism. **A** PBMCs were pre-activated with 10bpc fixed *E*. *coli* for 20 min, washed, and cultured for 6 hours in the absence (no N) or presence of equal numbers (1x N) of freshly isolated, or fixed neutrophils (n=9 for MAIT cells and n=7 for Vδ2^+^ T cells). **B** Pre-activated PBMCs were seeded in the lower compartment of a 0.4 μm pore size membrane transwell system and cultured for 6 hours in the absence (no N), presence of equal numbers (1x N), or triple the number (3x N) of freshly isolated neutrophils in the upper compartment. Control plates without transwells were run in parallel. TNFα and IFNγ expression was assessed by flow cytometry in MAIT (n=6) and Vδ2^+^ T cells (n=7). **C-D** Preactivated PBMCs were cultured for 6 hours in the absence (no N) or presence of equal numbers (1x N) of freshly isolated neutrophils and treated with **C** an isotype control (10 μg/mL), anti-CD11b blocking antibody (10 μg/mL), or catalase (4000 U/mL) (n=7), or **D** with L-arginine (5 mM) or nor-NOHA (500 μM) (n=6). TNFα expression by MAIT cells and Vδ2^+^ T cells was assessed by flow cytometry. **E** Untreated (UT) or pre-activated PBMCs (*E*. *coli* (p)) were cultured for 6 hours in the absence (no N) or presence of equal numbers (1x N) of freshly isolated neutrophils and treated with L-arginine (5 mM) or catalase (4000 U/mL) (n=8). Expression of CD3ζ by MAIT and Vδ2^+^ T cells was assessed by flow cytometry. All graphs show mean with SEM. Each data point represents an individual blood donor. **A**-**E** Repeated measures two-way ANOVA with Bonferroni’s multiple comparisons test was performed. * p<0.05, ** p<0.01, *** p<0.001, **** p<0.0001, ns = not significant.

Neutrophils have been shown to inhibit conventional T cells through a cell contact-dependent mechanism involving MAC-1 (CD18/CD11b) and the release of reactive oxygen species (ROS), including hydrogen peroxide (H_2_O_2_)^20^ We tested if this mechanism was responsible for the inhibitory effect seen on MAIT cells and Vδ2^+^ T cells. Blocking of MAC-1 with an antibody against CD11b did not prevent the inhibitory effect of neutrophils on TNFα and IFNγ production by MAIT cells and Vδ2^+^ T cells (Figure 3C, Supplementary Figure 2A). In contrast, removal of H O through catalase addition reversed the suppressive effect of neutrophils on MAIT cells and Vδ2^+^ T cells. Both TNFα and IFNγ production in samples treated with catalase were significantly higher than in control samples and were comparable to PBMCs alone for both cell types (Figure 3C, Supplementary Figure 2A). Catalase treatment of PBMCs alone also lead to significantly higher TNFα production in Vδ2^+^ T cells (Supplementary Figure 2A).

Neutrophils have further been described to inhibit T cell function through the release of arginase, resulting in the degradation of arginine, which is required for the expression of the CD3 ζ-chain (CD3ζ) on T cells.^26, 27, 28^ Thus, low levels of arginine can affect T cell activation. Supplementation with L-arginine partially reduced the inhibitory effect of neutrophils on MAIT cells and Vδ2^+^ T cells. TNFα expression was significantly higher than in control samples for both cell types (Figure 3D), though this effect was not consistently evident when production of IFNγ was assessed (Supplementary Figure 2D). TNFα production by Vδ2^+^ T cells in PBMCs alone was slightly increased with L-arginine treatment (Figure 3D). Inhibition of arginase with either nor-NOHA or L-valine did not reverse the suppressive effect of neutrophils on MAIT and Vδ2^+^ T cell activation (Figure 3D, Supplementary Figure 2B-D).

Both supplementation of L-arginine and addition of catalase prevented the inhibitory effect of neutrophils on CD107a upregulation on MAIT cells and Vδ2^+^ T cells (Supplementary Figure 3A). The effect of these treatments on granzyme B, perforin, and IL-17A expression was not as clear (Supplementary Figure 3B-D). A slight reduction of granzyme B expression in both cell types was seen in samples where neutrophils were present. This reduction was not seen in samples treated with catalase, however, the difference between non-catalase and catalase treated samples was not statistically significant (Supplementary Figure 3B).

CD3ζ expression on MAIT and Vδ2^+^ T cells was significantly reduced when neutrophils were present, even in untreated samples (Figure 3E). In contrast, CD3ε expression was significantly increased in the presence of neutrophils, and TCR expression was increased in MAIT cells but decreased in Vδ2^+^ T cells (Supplementary Figure 3E-F). L-arginine treatment did slightly increase CD3ζ expression, but not to the level of that in PBMCs alone. Addition of catalase did not significantly alter CD3ζ expression in either cell type. Interestingly, CD3ζ was expressed at higher levels by Vδ2^+^ T cells than MAIT cells, and in response to bacteria MAIT cells slightly downregulated CD3ζ, whereas it was slightly upregulated by Vδ2^+^ T cells (Figure 3E).

Another mechanism of T cell suppression is through adenosine. Binding of adenosine to adenosine receptors on T cells can inhibit or skew effector cytokine production. In iNKT cells it has been observed that adenosine inhibits IFNγ production but induces IL4 production.^29^ Neutrophils are known to be a source of extracellular adenosine,^30, 31, 32^ therefore, we tested if this might be a mechanism through which neutrophils inhibit MAIT cell and Vδ2^+^ T cell activation. Treatment of *E*. *coli* stimulated samples with the adenosine receptor antagonist, ZM241385, did not influence the effect of neutrophils on MAIT cell or Vδ2^+^ T cell activation (Supplementary Figure 4).

Overall, the inhibitory effect of neutrophils on MAIT cell and Vδ2^+^ T cell activation appeared to be dependent on cell contact, ROS but not MAC-1, partially on arginine concentration, and, for MAIT cells, on neutrophil viability.

### Activated MAIT cells kill neutrophils

Previously, CD3/CD28 bead activated MAIT cells have been reported to protect against neutrophil death.^22^ Here, we tested if ligand activated MAIT cells similarly promote neutrophil survival. Vα7.2^+^ (MAIT) cells were isolated and activated for either 4 or 24 hours using the MAIT cell ligand precursor, 5-A-RU, and adherent THP1 cells. 5-A-RU reacts non-enzymatically with methylglyoxal or glyoxal, present in metabolising cells, to form the MAIT cell activating ligand. Activated MAIT cells or conditioned media from the activation cultures were added to freshly isolated autologous neutrophils (Figure 4A). Neutrophil survival was measured by both Annexin V staining after 5 hours, as well as CD16 and viability staining after 24 hours. Adherent THP1 cells treated with 5-A-RU (control) or co-cultured untreated THP1 cells and MAIT cells (UT) were used as controls. Neutrophils treated with supernatant from either 4 or 24 hour activated MAIT cells showed poorer survival compared to control samples. The frequency of both Annexin V positive cells and dead cells were increased in samples treated with conditioned media from activated MAIT cells (Figure 4B+C). Neutrophils co-cultured with 4 hour pre-activated MAIT cells (at a ratio 1:5 MAIT cells to neutrophils) also showed higher levels of cell death, but not significantly higher Annexin V staining. This effect was dependent on the ratio of neutrophils to MAIT cells, with fewer MAIT cells (ratio 1:10 and 1:50 MAIT cells to neutrophils) resulting in less cell death, and returning to the level in control samples at the 1:50 ratio (Figure 4D+E). A reduction of Annexin V staining was also observed when neutrophils were treated with fewer MAIT cells (Figure 4D). In contrast, neutrophils treated with 24 hour activated MAIT cells showed no change in neutrophil viability and minimal change in Annexin V staining (Figure 4F+G). Treatment of neutrophils with 4 or 24 hour activated MAIT cells significantly reduced the frequency of CD16^high^ neutrophils; when fewer MAIT cells were added, the frequency of CD16^high^ neutrophils increased (supplementary figure 5A-B). Treatment with conditioned media from activated MAIT cells also lead to reduced frequencies of CD16^high^ neutrophils, although the differences were not statistically significant (supplementary figure 5C). Comparison of Annexin V and dead cell staining of neutrophils treated with conditioned media or cells from the controls (loosened adherent THP1 cells treated with 5-A-RU (control) and co-cultures of untreated adherent THP1 cells and MAIT cells (UT)) revealed a small reduction of Annexin V staining in samples treated with non-activated MAIT cells. No other treatment showed any significant effect on neutrophils (Supplementary Figure 5D-E).

**Figure 4:**
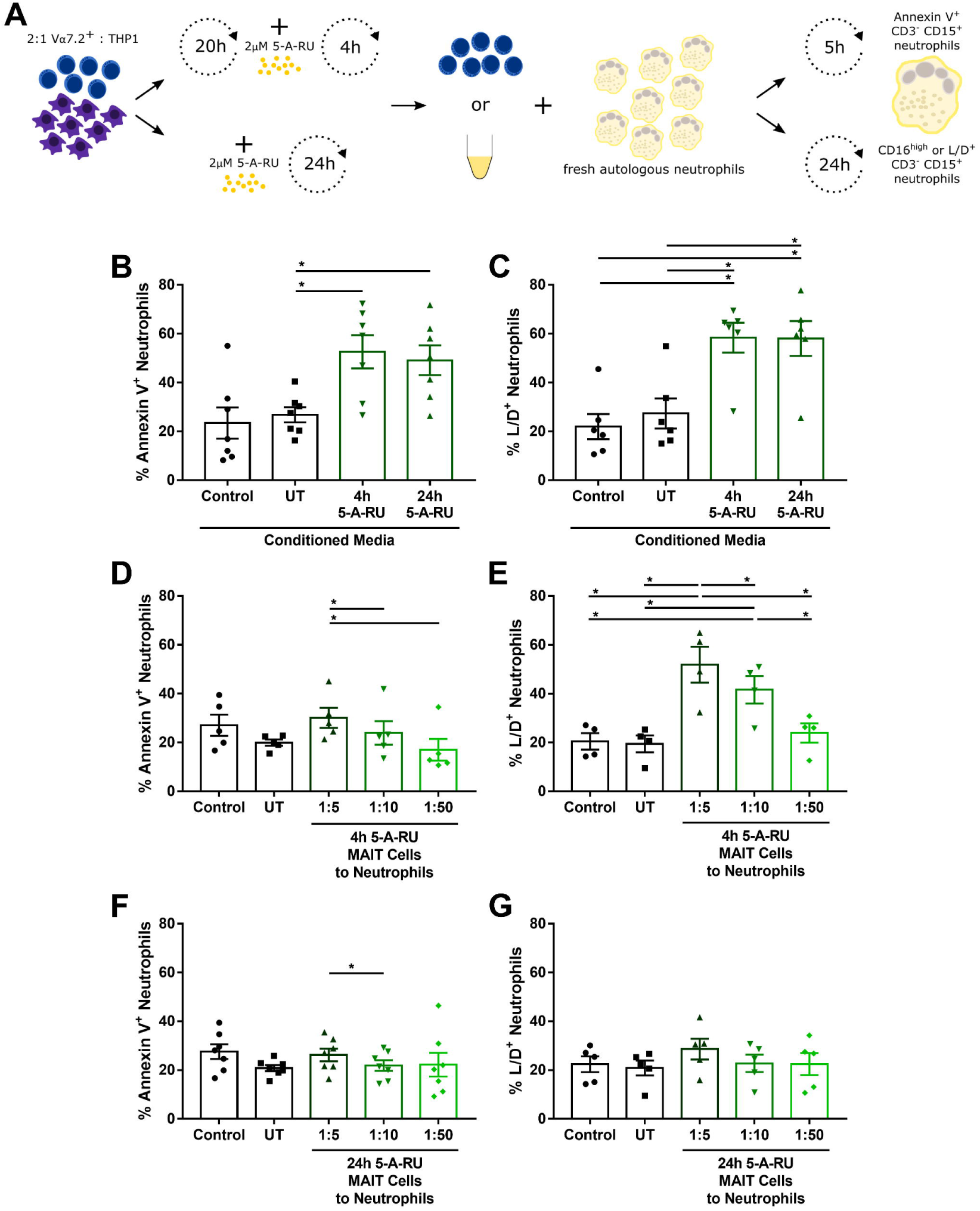
Activated MAIT cells kill neutrophils. **A** Isolated Vα7.2^+^ cells were seeded on adherent PMA-treated THP1 cells at a 2:1 ratio. Cells were either left untreated for 20 hours, then treated with 2 μM 5-A-RU for 4 hours, or treated with 2 μM 5-A-RU for 24 hours. Untreated co-cultures (UT) and THP1 cells treated with 2 μM 5-A-RU for 24 hours in the absence of Vα7.2^+^ cells (Control) were used as controls. Both non-adherent cells and conditioned media were harvested and co-cultured with freshly isolated autologous neutrophils. Annexin V staining (at 5 hours), dead cell staining, and CD16 expression (both at 24 hours) were assessed by flow cytometry. Neutrophils treated with conditioned media were stained for **B** Annexin V (at 5 hours) (n=7) and **C** dead cells (at 24 hours) (n=6). **D** Annexin V staining after 5 hours (n=5) and **E** dead cell staining after 24 hours of neutrophil co-culture with 4 hour 5-A-RU activated MAIT cells at different ratios, as indicated (n=4). **F** Annexin V staining after 5 hours (n=7) and **G** dead cell staining after 24 hours of neutrophil co-culture with 24 hour 5-A-RU activated MAIT cells at different ratios as indicated (n=5). **B**-**G** All graphs show mean with SEM. Each data point represents an individual blood donor. Repeated measures one-way ANOVA with Tukey’s multiple comparisons test was performed. * p<0.05.

### The amount of TNFα produced by MAIT cells influences neutrophil survival

Activated MAIT cells rapidly produce high levels of TNFα (see figure 2B).^8^ It has been described that high levels of TNFα (>10 ng/mL) induce TNFR dependent apoptosis in neutrophils ^33^. Therefore, we analysed whether the killing effect seen by conditioned media from cultures of activated MAIT cells was dependent on TNFα. Addition of a blocking antibody against TNFα significantly inhibited apoptosis and cell death in neutrophils treated with activated MAIT cell conditioned media. Viability staining after 24 hours of incubation was significantly reduced with addition of the TNFα blocking antibody, reducing it to the level of the controls (Figure 5A). Measurement of TNFα in the conditioned media of 4 and 24 hour 5-A-RU activated MAIT cells revealed mean concentrations of approximately 100 ng/mL and 70 ng/mL, respectively. Therefore, allowing for dilution, neutrophils were exposed to mean concentrations of >25 ng/mL and >17 ng/mL, respectively. In control samples, mean levels of TNFα between 1-10 ng/mL were measured (exposure of 0.25-2.5 ng/mL, respectively) (Figure 5B). Therefore, TNFα production by activated MAIT cells can induce neutrophil death.

**Figure 5:**
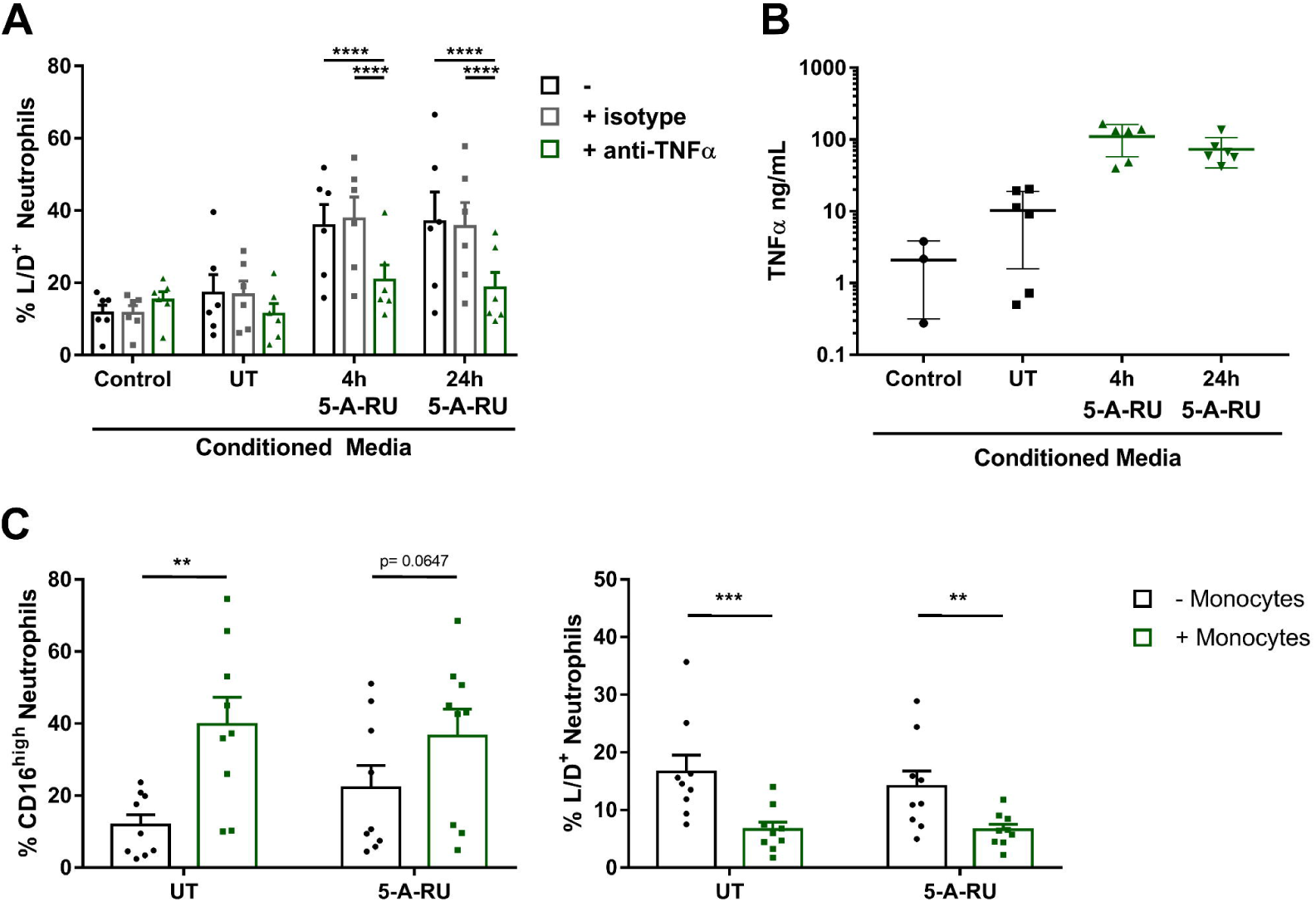
MAIT cell killing of neutrophils is TNFα dependent. **A** Dead cell staining of neutrophils treated with conditioned media from activated MAIT cells or controls (see Figure 4A) and isotype control or TNFα blocking antibody (both 10 μg/mL) for 24 hours, as indicated (n=6). **B** TNFα concentration of conditioned media from activated MAIT cells or controls (see Figure 4A) was assessed by ELISA, as indicated (n=3 for control; n=6 for all others). One THP1 cells only control was tested per experiment, each containing two MAIT cell donors. **C** CD16 and dead cell staining of untreated or 2 μM 5-A-RU treated neutrophils cultured with Vα7.2^+^ cells alone, or both with or without monocytes for 24 hours (n=9). **A**-**C** All graphs show mean with SEM. Each data point represents an individual blood donor. RM two-way ANOVA with Bonferroni’s multiple comparisons test was performed. ** p<0.01, *** p<0.001, **** p<0.0001.

## Discussion

Neutrophils are critical early responders to bacterial infection, phagocytosing and killing microbes.^15^ They also regulate the immune response by modulating other immune cells, including T cells.^34^ MAIT cells are also important early effector cells during bacterial infection and are present in mucosal tissues. Here, we have investigated the interaction between MAIT cells and neutrophils. We show for the first time that neutrophils suppress the activation of MAIT cells in response to bacteria. The ability of neutrophils to suppress MAIT cell activation was dependent on 1) neutrophil viability, 2) cell contact, 3) H_2_O_2_ but not MAC-1, and was independent of 4) arginase, but was weakened by arginine supplementation, and 5) was unaffected by adenosine receptor antagonist, ZM241385. We further investigated the effect of activated MAIT cells on neutrophil survival and found that, in contrast to CD3/CD28 bead activated MAIT cells,^22^ 5-A-RU activated MAIT cells produced TNFα at levels able to promote neutrophil death. Therefore, we propose that neutrophils and MAIT cells negatively regulate each other to control inflammation.

The effect of neutrophils on MAIT cell activation was investigated by assessment of a broad range of activation markers; comparison was made to another prominent innate-like T cell population, Vδ2^+^ T cells. IFNγ and TNFα production was substantially suppressed in both cell subsets, as was upregulation of CD69. In MAIT cells only, upregulation of 4-1BB and degranulation marker CD107a was also inhibited. Neutrophils did not have any significant effect on granzyme B, perforin, or IL-17A expression. Therefore, we can conclude that neutrophils suppress MAIT cell and Vδ2^+^ T cell activation and do not induce a major change in their effector phenotype. For MAIT cells, this suppression was seen both in co-culture of isolated cells as well as in whole PBMCs. The suppressive effect was maintained both when bacteria were pre-incubated with PBMCs and when MAIT cells were activated with 5-A-RU alone; this eliminated phagocytosis of bacteria by neutrophils as the cause of the suppression. In both co-culture and whole PBMC experiments, the presence of neutrophils had minimal effect on monocyte activation. Therefore, it is probable that neutrophils act directly on MAIT and Vδ2^+^ T cells. This is supported by the observation that neutrophils interact directly with γδ T cells.^19^ Moreover, neutrophils have been shown to inhibit conventional T cell activation through a cell contact-dependent mechanism involving MAC-1 (CD11b/CD18) and the localised release of H_2_O_2_in an immunological synapse.^20^ Suppression of MAIT cells and Vδ2^+^ T cells by neutrophils was dependent on H_2_O_2_, but not MAC-1, and cell contact was required. Neutrophils have similarly been implicated in the suppression of invariant natural killer cells (iNKT) through an unidentified cell contact-dependent mechanism.^18^ It is plausible that suppression of these two innate-like lymphocyte subsets is dependent on the interaction of other surface molecules which have either a direct inhibitory effect, such as PD-1,^35^ or facilitate cell binding, in addition to soluble factors, such as H_2_O_2_. Therefore, it is likely that neutrophils are able to affect activation of innate-like lymphocytes through a generalised mechanism dependent on cell contact, or proximity, and soluble factors.

Neutrophil viability was important for MAIT cell inhibition, however, the inhibition of Vδ2^+^ T cell activation was not dependent on neutrophil viability. Sabbione *et al*. previously showed that γδ T cell inhibition was dependent on viability.^19^ This contradiction may be explained by different experimental setups. Sabbione *et al*. used isolated γδ T cells and activated these using the Vδ2^+^ T cell ligand, (E)-4-Hydroxy-3-methyl-but-2-enyl pyrophosphate (HMB-PP), whereas we used whole PBMCs and a bacterial stimulus for activation.

The release of arginase by neutrophils and the resulting reduction in arginine concentration has previously been described to inhibit T cell activation through the downregulation of the CD3ζ-chain.^36, 37, 38^ Indeed, CD3ζ expression in both MAIT and Vδ2^+^ T cells was drastically reduced in the presence of neutrophils. L-arginine supplementation slightly increased CD3ζ expression in MAIT cells and Vδ2^+^ T cells, however, CD3ζ expression was still significantly lower than in PBMC samples alone. Therefore, the effect of neutrophils on CD3ζ expression cannot be explained by changes in L-arginine concentration alone. The potential of neutrophils to suppress MAIT cells and Vδ2^+^ T cells was reduced when L-arginine was added, however, inhibition of arginase using either L-valine or nor-NOHA did not have an effect. A previous report found arginine supplementation had no effect on neutrophil suppression of γδ T cells, though this was assessed using only one activation marker, CD25, and the experimental setup differed to this study.^19^ In addition to arginase activity, neutrophils are able to rapidly deplete exogenous arginine through importation and nitric oxide synthase (NOS) mediated metabolism to nitric oxide (NO) and citrulline.^39^ Future experiments should examine the effect of blocking NOS activity alone, as well as in combination with arginase inhibition, as these enzymes are known to compete for arginine.^40^

Previously, CD3/CD28 bead activated MAIT cells were shown to promote neutrophil survival, which was dependent on TNFα, IFNγ, and GM-CSF.^22^ The levels of TNFα produced in response to CD3/CD28 bead activation were, on average, over 1 ng/mL and less than 10 ng/mL. In our experimental setup, levels of TNFα were significantly higher, with 5-A-RU activated MAIT cells producing an average of over 100 ng/mL TNFα. High concentrations (>10 ng/mL) of TNFα have been shown to induce TNFR dependent apoptosis in neutrophils.^33^ Therefore, bead activation of MAIT cells results in a much milder activation of MAIT cells; not enough to produce sufficient TNFα to induce neutrophil death. TNFα levels in sputum of patients with acute respiratory distress syndrome and synovial fluid of patients with severe rheumatoid arthritis, have been shown to reach 10 ng/mL, though generally levels are lower than that. ^41, 42^ We recently found median levels of TNFα of 343 pg/mL (range 4 pg/mL – >7.0 ng/mL) in sputum samples of patients with pneumonia.^43^

We propose a model whereby activated MAIT cells recruit neutrophils to the site of infection^8^ and neutrophils suppress MAIT cells, preventing excess activation. At the same time, while moderately activated MAIT cells may support neutrophils survival, strongly activated MAIT cells will induce neutrophil apoptosis in a TNFα-dependent manner. Therefore, the balance between MAIT cells and neutrophils may be an important factor in providing an effective, but controlled immune response.

## Methods

### Cells

Peripheral blood mononuclear cells (PBMCs) were isolated from heparinised blood of healthy donors (collected with informed consent with approval from the University of Otago Human Ethics Committee (Health)) using Lymphoprep (Alere Technologies GmbH, Germany) and centrifugation at 800xg for 30min. PBMCs were used immediately or cryopreserved in foetal bovine serum (FBS) supplemented with 10% dimethyl sulfoxide (DMSO, Sigma Aldrich, USA) and stored in liquid nitrogen. Thawed PBMCs were rested overnight in R10 media (RPMI 1640 containing L-glutamine and supplemented with 1% Penicillin/Streptomycin and 10% FBS (all ThermoFisher Scientific, USA) at 37°C with 5% CO_2_. Neutrophils were isolated as previously using Lymphoprep separation.^44^ Vα7.2^+^ cells (referred to as MAIT cells) and CD14^+^ cells (referred to as CD14^+^ monocytes) were isolated using an anti-Vα7.2-PE labelled antibody (3C10; BioLegend, USA) and anti-PE microbeads or CD14 microbeads (Miltenyi Biotec, Germany), respectively, as per the manufacturer’s instructions. THP1 cells were cultured in R10 media. All experiments were performed in flat-bottom 96-well plates (ThermoFisher Scientific, USA), unless otherwise stated.

### Antibodies and Dyes

The following antibodies and labelled proteins were used: Vα7.2 PE (3C10), Vα7.2 PE-Cy7 (3C10), CD3 BV510 (OKT3), CD3 PE-Cy7 (UCHT1), CD14 BV605 (M5E2), CD15 Pacific Blue (W6D3), CD15 PE-Cy7 (W6D3), CD16 BV510 (3G8), CD69 FITC (FN50), CD80 FITC (2D10), CD86 APC (IT2.2), CD107a PE (H4A3), CD137 (4-1BB) BV421 (4B4-1), CD247 (CD3ζ) FITC (6B10.2), granzyme B FITC (QA16A02), HLA-DR AF700 (LN3), IFNγ PerCP-Cy5.5 (4S.B3), IL-17A PE/Dazzle 594 (BL168), MR1 PE (26.5), Perforin PerCP-Cy5.5 (B-D48), TNFα FITC (MAb11), Vδ2 Pacific Blue (B6), Vδ2 PerCP (B6), Annexin V AF647 (Annexin A5) (all BioLegend, USA), CD161 APC (191B8, Miltenyi Biotec, Germany). Samples were stained with Live/Dead fixable near IR dye (ThermoFisher Scientific, USA).

### Inhibitors, Cytokines and Functional Antibodies

The following antibodies were used for blocking experiments: anti-IL-12 (Miltenyi Biotec, Germany), anti-MR1 (26.5), anti-CD11b (ICRF44), anti-TNFα (MAb11), and IgG1 (MOPC-21) and IgG2a (MOPC-173) isotype controls (all BioLegend, USA). Catalase from *Corynebacterium glutamicum*, L-arginine, L-valine, and ZM241385 were purchased from Sigma-Aldrich (USA), and nor-N^ω^-hydroxy-L-arginine (nor-NOHA) from Cayman Chemical (USA).

### Bacteria

*E*. *coli* (HB101) were grown overnight in Luria Broth (ThermoFisher Scientific, USA). For fixation, *E*. *coli* were washed twice in phosphate buffered saline (PBS), fixed for 20 min at room temperature in 2% paraformaldehyde/PBS (PFA), and washed twice in PBS. Bacterial concentration was estimated by flow cytometry using 123count eBeads (ThermoFisher Scientific, USA). Ten bacteria per antigen presenting cell/PBMC (bpc) were used, unless otherwise stated.

### 5-A-RU

5-A-RU was synthesised as described previously.^8^

### MAIT cell activation

10^5^ Vα7.2^+^ cells and 5×10^5^ neutrophils (1:5 ratio) were co-cultured for 6 or 24 hours in a flat-bottom 96-well plate and treated with *E*. *coli* with or without blocking antibodies against MR1, IL-12, or isotype controls. Alternatively, 10^5^ Vα7.2^+^ cells were co-cultured for 6 or 24 hours with or without 10^5^ CD14^+^ cells and/or 5×10^5^ neutrophils (1:1:5 ratio) and treated with *E*. *coli* or 2 μM 5-A-RU. In some cases, neutrophils were pre-treated with *E*. *coli* for 30 min, washed, then added to the co-cultures. For experiments with PBMCs, 10^6^ cells were mixed with 10^6^ neutrophils (1:1 ratio) and treated with *E*. *coli*. In some cases, PBMCs were pre-treated with *E*. *coli* for 20 min, washed twice, then mixed with neutrophils. For analysis of MR1 expression, the antibody was mixed with 20% mouse serum (Sigma-Aldrich, USA) in PBS and added at the start of treatment.^45^ For analysis of CD107a, the antibody was added at the start of treatment. Brefeldin A (3 μg/mL, BioLegend, USA) was added for the final 4 hours of culture for the analysis of cytokine expression.

### Analysis of mechanism of MAIT cell suppression

Neutrophils were fixed for 10 min in 2% PFA, washed twice in PBS, and resuspended in R10. Co-cultures were treated with 4000 U/mL catalase, 10 μg/mL blocking antibody against CD11b or an isotype control, 5 mM L-arginine, 20 mM L-valine, 500 μM nor-NOHA, or 1 μM ZM241385 (A2AR antagonist) where indicated. For analysis of cytokine expression, brefeldin A (3 μg/mL, BioLegend, USA) was added for the final 4h of culture. Transwell (0.4 μm pore size, polycarbonate membrane; Corning, USA) experiments were performed by addition of 3.5×10^6^ PBMCs to the bottom well and 3.5-10.5×10^6^ neutrophils to the top well; appropriate controls were performed alongside in 24-well plates.

### Effect of activated MAIT cells on neutrophils

10^5^ THP1 cells were treated with 20 ng/mL phorbol myristate acetate (PMA, Sigma Aldrich, USA) for 24 hours, washed, and rested for 24 hours, to generate adherent monocyte-derived macrophages. 2×10^5^ MAIT (Vα7.2^+^) cells were then activated on these adherent THP1 cells for 4 or 24 hours with 2 μM 5-A-RU. Controls containing untreated MAIT cells on THP1 cells or 2 μM 5-A-RU treated THP1 cells without MAIT cells were prepared in parallel. Following, neutrophils were isolated from fresh blood of the same donor. 0.5-5×10^4^ of 2 μM 5-A-RU-activated Vα7.2^+^ (MAIT) cells, 5×10^4^ untreated MAIT cells, 2 μM 5-A-RU treated THP1 cells (cells that dislodged during pipetting), or 50 µL of their supernatant were mixed with 2.5×10^5^ neutrophils (200 µL total) and incubated for 5 or 24 hours. In some experiments, 10 μg/mL blocking antibody against TNFα or an isotype control were added.

### Immunostaining

Samples were stained with various panels of antibodies against CD3, CD14, CD15, CD16, CD69, CD80, CD86, HLA-DR, CD137 (4-1BB), CD161, and Vδ2, as well as a Live/Dead fixable near IR dye for 25 min at 4°C, washed and fixed for 10 min in 2% PFA at room temperature. For intracellular protein analysis, samples were washed in PBS and 1X Permeabilization Wash Buffer (BioLegend, USA), prior to staining for CD3, CD247 (CD3ζ), Vα7.2, granzyme B, IFNγ, IL-17a, perforin, and TNFα for 25 min at room temperature.

For Annexin V staining, cells were stained with antibodies against CD3, CD15, and CD16 and with Live/Dead fixable near IR dye for 25 min on ice, washed once in PBS and once in annexin binding buffer (10 mM HEPES (pH 7.4), 140 mM sodium chloride, 2.5 mM calcium chloride; all Sigma-Aldrich, USA) prior to staining with Annexin V for 10 min on ice. Samples were kept on ice and acquired immediately on the flow cytometer.

### Flow Cytometry

Flow cytometry was performed on a BD FACSCanto II or a BD Fortessa (BD Bioscience, USA). Gating strategies for different panels and experiments are shown in Supplementary Figures 6-7. Analysis was performed with FlowJo version 10.4 (TreeStar, USA).

### TNFα ELISA

Conditioned media was analysed for TNFα concentration using the human uncoated ELISA kit with plates from ThermoFisher Scientific (USA). Manufacturer’s instructions were followed and plates analysed using the Varioskan Lux plate reader (ThermoFisher Scientific, USA). A second order polynomial (quadratic) fitted standard curve was used to interpolate sample concentrations using GraphPad prism software, version 7.00 (USA).

### Statistical Analysis

Data were analysed in GraphPad Prism software version 7.00. Unless otherwise stated, means with the standard error of the mean (SEM) and all data points are shown. Specific statistical tests used are described in the figure legends. Generally, two-tailed, repeated measures one or two-way ANOVA were used with multiple comparisons tests where appropriate. A p value below 0.05 was considered statistically significant.

## Acknowledgements

This work was supported by the Health Research Council (HRC) of New Zealand.

## Author contributions

M.S., R.F.H., R.L. performed the experiments. M.S. analysed the data. M.S., A.K., J.E.U. designed the experiments. M.S., J.E.U. managed the study. S.d.l.H., J.T., A.V., synthesised the 5-A-RU. M.S., J.E.U. conceived the work and wrote the manuscript which was revised and approved by all authors.

## Competing interests

All authors declare no competing interests.

---

**Supplementary Figure 1**: **Neutrophils are poor activators of MAIT cells**. Vα7.2^+^ cells and neutrophils (1:5 ratio) were activated with 10bpc fixed **E**. **coli** and incubated **A** ± blocking antibodies against MR1 and/or IL-12 for 22 hours, or **B-D** ± CD14^+^ monocytes (1:1 ratio to Vα7.2^+^ cells) for 6 hours, with brefeldin A added for the final 4 hours. **E** PBMCs were pre-incubated (p) with 10bpc fixed *E*. *coli* for 20 min prior to washing, addition to neutrophils, and incubation for 6 hours, with brefeldin A added for the final 4 hours. **A** IFNγ (n=7) and **B-C** IFNγ and TNFα (n=2) production in MAIT cells (CD161^++^Vα7.2^+^CD3^+^) is shown. **D-E** geometric mean of fluorescence intensity of TNFα in CD14^+^ monocytes is shown for (n=8). Mean with SEM of IFNγ and/or TNFα production in MAIT cells (CD161^++^Vα7.2^+^CD3^+^) or CD14^+^ monocytes is shown. Each data point represents an individual blood donor. **A** Repeated measures one-way ANOVA with Tukey multiple comparisons test **B-D** Repeated measures one-way ANOVA with Bonferroni’s multiple comparison test of pre-selected pairs was performed. **E** Repeated measures two-way ANOVA with Bonferroni’s multiple comparisons test. ** p<0.01, *** p<0.001, ns = not significant.

**Supplementary Figure 2**: **Mechanism of MAIT cell suppression by neutrophils**. Pre-activated PBMCs were cultured for 6 hours in the absence (no N) or presence of equal numbers (1x N) of freshly isolated neutrophils and treated with **A** an isotype control (10 μg/mL), anti-CD11b blocking antibody (10 μg/mL), or catalase (4000 U/mL) (n=7), or **B** L-arginine (5 mM) or nor-NOHA (500 μM) (n=6), or **C**-**D** L-arginine (5 mM) or L-valine (20 mM) (n=7). IFNγ (**A**, **B**, and **D**) and TNFα (**C**) expression was assessed by flow cytometry in MAIT cells and Vδ2^+^ T cells. **A**-**D** All graphs show mean with SEM. Each data point represents an individual blood donor. Repeated measures two-way ANOVA with Bonferroni’s multiple comparisons test was performed. * p<0.05, ** p<0.01, *** p<0.001, **** p<0.0001.

**Supplementary Figure 3**: **Suppression of MAIT cells by neutrophils**. **A-F** Pre-activated PBMCs were cultured for 6 hours in the absence (no N) or presence of equal numbers (1x N) of freshly isolated neutrophils and treated with L-arginine (5 mM) or catalase (4000 U/mL). Expression of **A** CD107a (n=7), **B** granzyme B (n=5), **C** perforin (n=5), **D** IL-17A (n=5), **E** CD3ε (n=6), and **F** Vα7.2 and Vδ2 (respectively) (n=8) was assessed by flow cytometry on MAIT cells and Vδ2^+^ T cells. **A**-**F** All graphs show mean with SEM. Each data point represents an individual blood donor. Repeated measures two-way ANOVA with Bonferroni’s multiple comparisons test was performed. * p<0.05, ** p<0.01, *** p<0.001, **** p<0.0001, ns = not significant.

**Supplementary Figure 4: Effect of adenosine on MAIT cell activation**. **A-B** Pre-activated PBMCs were cultured for 6 hours in the absence (no N) or presence of equal numbers (1x N) of freshly isolated neutrophils and treated with ZM241385 (1 μM) (n=6). TNFα (**A**) and IFNγ (**B**) expression was assessed by flow cytometry in MAIT cells and Vδ2^+^ T cells. All graphs show mean with SEM. Each data point represents an individual blood donor. Repeated measures two-way ANOVA with Bonferroni’s multiple comparisons test was performed.

**Supplementary Figure 5: Effect of activated MAIT cells on neutrophil viability**. **A**-**C** CD16 staining of neutrophils after treatment for 24 hours with **A** 4 hour (n=6), or **B** 24 hour (n=8) 5-A-RU activated MAIT cells at different ratios, as indicated, or **C** treated with conditioned media as indicated (n=7). **D** Annexin V staining after 5 hours (n=7), or **E** dead cell staining after 24 hours (n=5) of untreated neutrophils (-), co-cultured with dislodged control THP1 cells or untreated MAIT cells, or conditioned media, as indicated. **A**-**D** All graphs show mean with SEM. Each data point represents an individual blood donor. Repeated measures one-way ANOVA with Tukey’s multiple comparisons test was performed. * p<0.05, ** p<0.01.

**Supplementary Figure 6: Gating strategy for identification of MAIT cells**, **Vδ2 cells**, **and CD14 monocytes**. All cells were gated for single cells, prior to separation into monocyte and lymphocyte subsets. Cells were further gated for viable cells, identified by live/dead cell staining. CD14^+^ monocytes are defined as CD14^+^CD15^-^ cells on the monocyte subset. MAIT cells are identified as CD14^-^CD15^-^CD3^+^CD161^++^Vα7.2^+^ cells and Vδ2^+^ cells are identified as CD14^-^CD15^-^CD3^+^Vδ2^+^ cells, each on the lymphocyte subset.

**Supplementary Figure 7: Gating strategy for identification of neutrophils**. All cells were gated for single cells, prior to the identification of neutrophils by forward and side scatter. Neutrophils were further defined as CD15^+^ cells. Gating strategies for additional markers, CD16^high^ and Annexin V, are shown. For each cell subset, dead cells were excluded from analysis by live/dead staining.

